# Thermal sensitivity of lizard embryos indicates a mismatch between oxygen supply and demand at near-lethal temperatures

**DOI:** 10.1101/788687

**Authors:** Joshua M. Hall, Daniel A. Warner

## Abstract

Aspects of global change (e.g. urbanization, climate change) result in novel, stressful thermal environments that threaten biodiversity. Though much research quantifies the thermal sensitivity of adult organisms, effects of global change on developing offspring (e.g. embryos) are also important. Oviparous, non-avian reptiles have received considerable attention because eggs are left to develop under prevailing environmental conditions, making them vulnerable to increases in ambient temperature. Though many studies assess embryo thermal tolerance and physiology in response to long-term (i.e. chronic), constant incubation temperatures, fewer assess responses to acute exposures which are more ecologically relevant for many species. We subjected eggs of the brown anole lizard (*Anolis sagrei*) to heat shocks, thermal ramps, and extreme diurnal fluctuations to determine the lethal temperature of embryos, measure the thermal sensitivity of embryo heart rate and metabolism, and quantify the effects of sub-lethal but stressful temperatures on embryo development and hatchling phenotypes and survival. Most embryos died at heat shocks of 45 or 46 °C, which is ∼12 °C warmer than the highest constant temperatures suitable for development. Heart rate and O_2_ consumption increased with temperature; however, as embryos approached the lethal temperature, heart rate and CO_2_ production continued rising while O_2_ consumption plateaued. These data indicate a mismatch between oxygen supply and demand at high temperatures. Exposure to extreme, diurnal temperature fluctuations depressed embryo developmental rates and heart rates, and resulted in hatchlings with smaller body size, reduced growth rates, and lower survival in the laboratory. Thus, even brief exposure to extreme temperatures can have important effects on embryo development, and our study highlights the role of both immediate and cumulative effects of high temperatures on egg survival. Such effects must be considered to predict how populations will respond to global change.

## 1. INTRODUCTION

Multiple aspects of global change (e.g. urbanization, climate change) create novel, stressful thermal environments that threaten biodiversity across the planet (McDonald et al., 2008; Sinervo et al., 2010). Much research on this topic quantifies how adult organisms respond to high temperatures (e.g. Sinervo et al., 2010; Battles and Kolbe, 2019); however, the effect of global change on developing offspring (e.g. embryos) is less studied, but also important (Ma et al., 2018; Burggren, 2018). Embryos are particularly sensitive to thermal stress due to a relatively narrow thermal tolerance compared to adults (Pörtner et al. 2017) and little to no capacity to behaviorally thermoregulate (Cordero et al. 2018; but see Du and Shine 2015). Additionally, because many important, thermally sensitive processes occur during development (e.g. organogenesis), thermal stress at this stage can induce life-long negative effects (Shine et al., 2005; Kaiser et al., 2016). Finally, egg mortality can drive population cycles (Chalcraft and Andrews 1999); therefore, the thermal sensitivity of embryos can influence species distributions and population persistence in the face of global change (Ma et al., 2018; Carlo et al. 2018). Thus, to understand how biodiversity will respond to novel thermal conditions, it is critical to quantify embryo responses to extreme thermal variation in natural habitats (Burggren, 2018).

Non-avian reptiles (henceforth, “reptiles”) have contributed greatly to our understanding of embryo thermal ecology (Noble et al., 2018; Refsnider et al., 2019), and many species are threatened by global change (Sinervo et al., 2010; Santidrián Tomillo et al., 2015). A recent flurry of work has synthesized existing egg incubation studies (While et al., 2018; Warner et al., 2018; Booth 2018; Noble et al. 2018), demonstrating that most researchers use a series of constant incubation temperatures to define the upper thermal limits for development (Andrews and Schwarzkopf 2012). Constant temperatures may be appropriate for species that construct relatively deep nests that experience little thermal variation; however, most eggs incubate in nests that exhibit daily fluctuations in temperature (Booth 2018). The effects of fluctuating temperatures differ widely from those of constant temperatures for a diversity of phenotypes (Bowden et al., 2014; Warner and Shine 2011; Noble et al., 2018); therefore, constant temperature treatments are insufficient to assess the effects of thermal stress on many wild populations. Moreover, most studies have used incubation treatments that persist throughout development (i.e. chronic exposure), but we know very little about the immediate or cumulative effects of brief (i.e. acute) exposure(s) to stressful temperatures (Angilletta et al., 2013). This knowledge gap should be filled for two reasons. First, it limits our ability to conduct broad, comparative analyses of embryo responses to ecologically relevant incubation temperatures. Indeed, such analyses currently depend on constant temperature incubation studies and ignore the effects of acute exposure to thermal extremes (e.g. Andrews and Schwarzkopf 2012). Second, in some contexts, maximum nest temperatures can drive the evolution of life-history traits more so than mean temperatures (Shine et al., 2003). Given that nest temperatures often fluctuate above the critical thermal maximum for some species (e.g. Angilletta et al., 2013; Sanger et al., 2018) and that global change will cause nest temperatures to rise in both mean and variance, more studies of acute exposure to thermal stress are needed.

The brown anole lizard (*Anolis sagrei*), is an excellent model for studies of environmental variability and development (Hall et al., 2019). Protocols for their captive husbandry are established (Sanger et al., 2008a), they are relatively fecund in captivity allowing for robust sample sizes (Hall et al., 2018), and their developmental staging series has been described (Sanger et al., 2008b). Females construct shallow nests across a diversity of habitats; thus, in the wild, embryos experience relatively large thermal variation during incubation (Sanger et al., 2018; Tiatragul et al., 2019) and temperature has important effects on embryo development, egg survival, and hatchling phenotypes (Pearson and Warner 2018). Moreover, daily fluctuations in nest temperature often reach extremely warm temperatures (> 40 °C, Sanger et al., 2018); indicating that embryos might have physiological mechanisms for ameliorating the adverse effects of acute exposure to high temperatures. Thus, the brown anole can serve as an excellent model to study the effects of extreme thermal variation on development.

We used eggs of the brown anole to quantify the immediate and cumulative effects of acute thermal stress on development and obtain novel, baseline information on physiology and thermal tolerance for this species. Our objectives were to 1) determine the acute lethal temperature for *A. sagrei* embryos, 2) quantify the thermal sensitivity of embryo physiology across nest temperatures and near-lethal temperatures 3) and assess the effects of repeated exposure to sublethal, but stressful temperatures on embryo development and hatchling phenotypes and survival. To determine the acute lethal temperature (T_LETHAL_) for embryos, we exposed eggs to 1-hour heat shocks of increasingly high temperatures. To quantify embryo physiology, we measured embryo heart rates and oxygen consumption (henceforth VO_2_) across temperatures from 22 to 47 °C, which encompasses the complete range of nest temperatures recorded for this species (Sanger et al., 2018; Tiatragul et al., 2019). To understand the effect of repeated exposures to sublethal temperatures, we subjected eggs to four exposures of an extreme fluctuation in nest temperature and measured aspects of embryo physiology, hatchling morphology, growth, and survival in the laboratory. Understanding the relationship between thermal stress and embryo development is vital to predict how populations will respond to global change.

## 2. METHODS

### 2.1 Determining T_LETHAL_ via heat shock

We collected adult lizards (n=60 females and n=12 males) from Palm Coast, FL (coordinates: 29.602199, −81.196211) from 22 to 24 March 2019. Lizards were transported to Auburn University and housed in screen cages (45 x 45 x 92 cm; Repti-Breeze, Zoo Med Inc.) in a 5:1 female:male ratio, and maintained at 27 °C with a 12:12 hr light:dark cycle using Reptisun 5.0 UVB bulbs (Zoo Med Inc.) and plant grow bulbs (model F40; General Electric Co.). We fed lizards 3 crickets each, dusted with vitamins and calcium twice per week and misted cages with water daily. Nest pots were provided with a mixture of peat moss and potting soil. We collected eggs twice per week from 7 to 20 June and placed all eggs in 60 mm petri dishes half-filled with moist vermiculite (−150 kPa) and wrapped with parafilm to prevent desiccation. Eggs were incubated at temperatures that fluctuated in a daily sine wave with amplitude of 2.4 °C and a mean of 26.3 °C, which is similar to nest temperatures at our field site (see Pearson and Warner 2018). We subjected all eggs (n=72) to a series of 1-hour heat shocks starting at 44 °C. Eggs that survived were given 3-4 days to recover and were heat-shocked at 45 °C. We repeated this process, increasing the heat shock by 1 °C, until all eggs were dead. For each egg, we recorded the temperature at which it died (i.e. lethal temperature; T_LETHAL_). Due to variation in egg-laying date among females, eggs ranged in age from 4 to 17 days post oviposition at the time of treatment.

To apply heat shocks, we placed eggs in glass jars (59mL FLINT s/s) that were ¾ filled with moist vermiculite (−150 kPa) and covered with half of a 60 mm petri dish to reduce water loss but allow gas exchange. After the heat shock, we checked each egg for a heartbeat using the Buddy^®^ heart rate monitor (Hulbert et al., 2017). Eggs with no heartbeat were considered dead and were returned to the fluctuating incubator and monitored daily for additional signs of mortality (e.g. fungal growth). All eggs without a heartbeat eventually shriveled and molded; thus, this method was effective.

To estimate T_LETHAL_ while controlling for variation in egg size and age, we performed a linear regression with T_LETHAL_ as the response variable and oviposition date (converted to Julian day of the year) and egg mass (which was measured prior to treatment) as fixed effects. Egg mass and age were centered at zero prior to analysis.

### 2.2 Thermal sensitivity of embryo physiology

We collected additional eggs from a different breeding colony (n=38 female; n=12 males) that was captured from 18-19 March 2018 from Pinecrest, FL (coordinates: 25.678125, −80.287655). Husbandry was as previously described; however, we housed females individually in cages (29 × 26 × 39 cm; height × width × depth) and rotated males among females. We used 11 randomly selected eggs collected 25 May to 1 June 2018 to determine the thermal sensitivity of embryo heart rate. Eggs were incubated in petri dishes (as previously described) at a constant 28 °C for 1 week prior to measurements. Thus, embryos varied in age from 7-14 days post oviposition; however, embryo heart rate does not covary with age in the first few weeks of development (Hulbert et al., 2017). Heart rate measures occurred over the course of 3 days. On day 1, eggs were kept at room temperature (∼ 22 °C) for 24 hours. On day 2, eggs were slowly (3°C per hour) raised from 22 to 39 °C inside a Memmert brand IPP 55 Plus incubator. We programmed the incubator to stop increasing temperature for approximately ½ hour at various target temperatures (22, 26, 29, 31, 34, 37, and 39 °C). During these intervals, we measured heart rates. Eggs were quickly removed (one at a time) from the incubator and placed in the heart rate monitor which was housed in another incubator set to the target temperature. Eggs remained in the monitor for 45 to 60 s before we recorded a heart rate. A thermocouple was secured to the inside of the monitor, and we recorded the air temperature inside the monitor with each heart rate. After heart rate measurements, eggs were returned to the Memmert incubator to increase to the next target temperature. On day 2, using the same eggs, we measured heart rates from 40 to 47 °C. Heart rates were measured at 40 °C (after bringing them to 40 °C by 3 °C per hour), and we increased the temperature of eggs at a steady rate (∼ 3 °C per hour) while measuring heart rate periodically (at approximately 1 °C intervals) until each egg was dead (i.e. no heart rate).

A different subset of eggs was used to determine the thermal sensitivity of embryo VO_2_ (n=45 eggs; i.e. all eggs collected from 23 – 30 July 2018). These eggs were incubated in petri dishes (as previously described) at a constant 28 °C for 1 week prior to measurements. Thus, embryos were between 7-14 days since oviposition at time of measurement. These eggs were randomly allocated to 1 of 9 temperature treatments (21, 25, 29, 33, 35, 39, 41, 44, 47 °C; n=5 per treatment). We measured the VO_2_ of eggs while they incubated at their assigned temperature over a 30-minute period using a Qubit Q-box RP1LP respirometer (Qubit Biology Inc., Kingston, ON) via dynamic injection. After measurements, each egg was placed in the heart rate monitor to determine survival. See Supporting information S1 for details about measuring VO_2_. CO_2_ production was simultaneously measured so we could calculate respiratory quotients (CO_2_ produced/O_2_ consumed, i.e. RQ) at each temperature.

To determine the thermal sensitivity of heart rate, we used the temperatures recorded inside the heart rate monitor at the time each heart rate was measured (rather than the nominal target temperature). We performed two linear mixed effects models with heart rate as the response variable and temperature as the independent variable. One model assumed the relationship was linear and the other assumed it was a second-degree polynomial (i.e. linear plus quadratic term). Egg ID was a random effect in all models. Best fit models were determined with likelihood ratio tests. Two data points were removed from the analysis because they were extreme outliers and likely represent eggs near death (see results).

To analyse VO_2_, we performed two general linear models: one was an asymptotic model, and the other was a second-degree polynomial. The best fit model was determined with a likelihood ratio test. We excluded eggs incubated at 47 °C because they died during treatment which prevents us from reliably calculating VO_2_. Preliminary analysis revealed no relationship between egg mass and VO_2_ (p=0.66) so we omitted egg mass from the model.

### 2.3 Repeated exposure to sublethal temperatures

On 4 March 2017, adult lizards (n=30 females; n=15 males) were captured from Pinecrest, FL and housed as described in section 2.2 above. We collected eggs three times per week and used all eggs produced from 3 to 17 July (n= 71). For each egg, we recorded the mass, date of oviposition, and maternal identity. Eggs were individually placed in petri dishes as previously described and randomly assigned to one of two incubation treatments (described below) and placed in an incubator that repeated a daily thermal fluctuation that was suitable for successful development and created from field nest temperatures (Tiatragul et al., 2017; Fig S1). Because anoles lay a single egg every one or two weeks, we randomly assigned each female’s first egg to a treatment and alternated subsequent eggs between the two treatments. Treatments subjected eggs to either zero (i.e. control) or four (i.e. experimental) exposures to an extreme fluctuation in temperature (henceforth, “thermal spike”) measured from the field (see Hall and Warner 2018). The peak temperature of this thermal spike was 43 °C. We chose 4 exposures to this peak temperature because a previous study showed that 1 or 2 exposures reduce egg survival and hatchling body size (Hall and Warner in review). Hatching success was 82 and 78% for 1 and 2 spikes, respectively, compared to 93% for controls. Hatchling SVL was reduced by 0.5 and 1% compared to controls for 1 and 2 spikes, respectively. Body mass was reduced by 1.6 and 2.5% compared to controls for 1 and 2 spikes, respectively. Though not statistically clear (p > 0.05), hatching success and body size declined with an increase in the number of exposures, suggesting that additional exposures (i.e. 4) may have a larger effect. Because eggs were collected over a 2-week period, eggs varied in age at the time of treatment. Figure 1 provides an overview of the experimental design and Figure S2 provides more details of temperature treatments.

**Figure 1.**
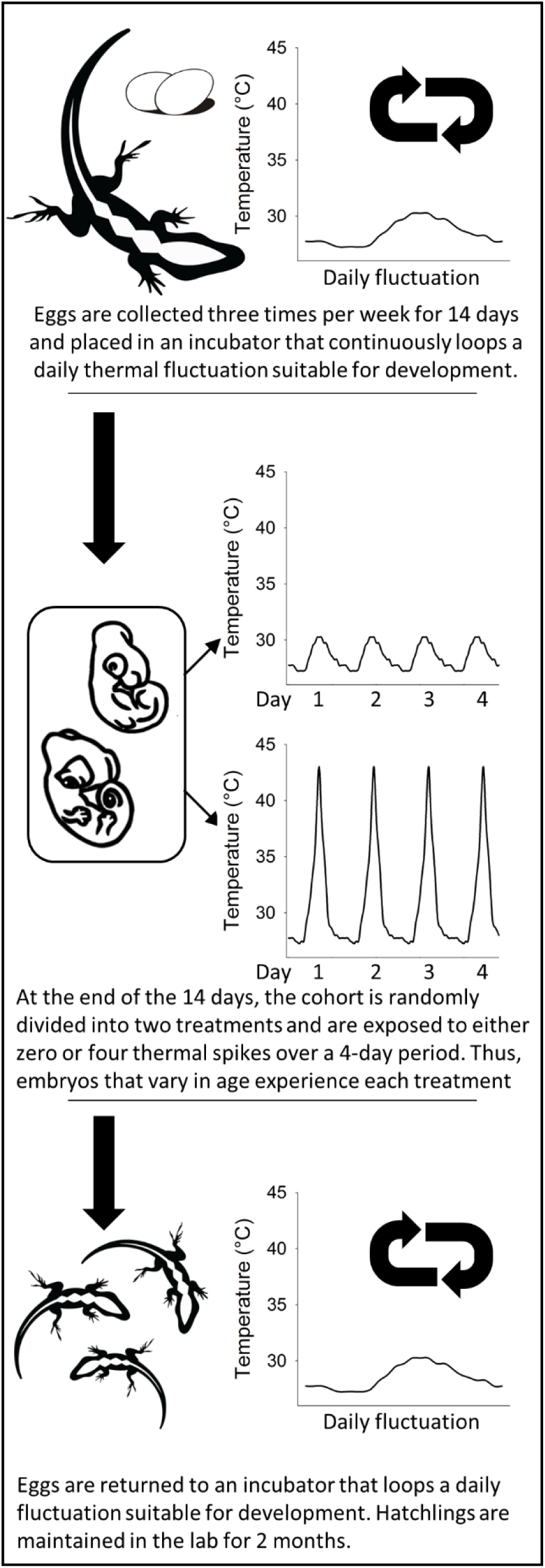
Overview of experimental design to assess repeated exposure to sublethal temperatures.

To quantify latent effects of thermal spikes on embryo physiology, we measured embryo heart rates using the Buddy^®^ egg monitoring system as previously described. For each egg, we measured heart rate at 28 °C on the day before and on each of two days after treatments.

For all hatchlings (n=61), we measured snout-vent length (SVL; to nearest 0.01 mm) and tail length (to nearest 0.01 mm) with a digital calliper, and hatchling mass (to nearest 0.0001 g). We kept hatchlings in cages that were identical to those described for adults in section 2.2. We aimed to keep 6 hatchlings per cage (3 from each treatment); however, due to differences in egg survival per treatment (see results) there were n=1 cage with 5 lizards and n=8 cages with 6 lizards. To minimize large age discrepancies among cage-mates, we filled cages in the order that lizards hatched; thus, the last 8 control lizards that hatched were not assigned to a cage but were euthanized because there were no more experimental hatchlings to serve as cage mates. We fed hatchlings fruit flies, *ad lib*, dusted with vitamins and calcium and misted cages with water each day. Once hatchlings reached approximately 2 months of age, we measured the final SVL and body mass of all survivors.

To assess the effects of thermal spikes on developmental rates, hatchling morphology (SVL, mass, tail length), and hatchling growth rates, we performed linear mixed effects models (LMMs) with initial mass (egg mass at oviposition for developmental rate and hatchling morphology; body mass at hatching for hatchling growth), embryo age at time of treatment, treatment (0 vs 4 spikes), and an age by treatment interaction as fixed effects. Maternal ID was the random effect for all analyses except cage ID was the random effect for hatchling growth rates. The interaction term was not statistically significant in any model, so it was omitted (all p-values > 0.13). Embryo age was the first day of treatment minus the day the egg was collected. To convert incubation periods to developmental rates, we divided the number of stages that embryos traverse from oviposition to hatching (15, Sanger et al. 2008b) by the total number of days from egg collection to hatching. Hatchling growth rate was the final mass of each hatchling minus its body mass at hatching and divided by the total number of days it was in captivity.

We analyzed egg and hatchling survival with generalized linear mixed effects models (GLMMs) with maternal ID as a random effect for egg survival and cage ID as a random effect for hatchling survival. We assumed a binomial distribution. Due to convergence issues, our final model for egg survival included only treatment as a fixed effect and no random effect. Only one egg died in the control group; so, we split the data by treatment and analyzed the experimental group with embryo age and egg mass as fixed effects to determine how survival might change with embryo age. For hatchling survival, we removed the random effect (i.e. hatchling cage) to achieve model convergence; however, raw survival estimates suggest some variation in survival among cages (6 cages = 0.50, 1 cage =0.67, 2 cages =0.83).

To analyze heart rates before and after thermal spikes, we performed a LMM with heart rate as the dependent variable and day (day before treatment, one day post-treatment, or two days post-treatment), treatment (0 vs 4 spikes), and a treatment by day interaction as fixed effects. The exact temperature in the heart rate monitor and embryo age were covariates.

All data analyses were performed in R (ver. 3.5.1; R Core Team 2018). For GLMMs, we used the “glmer” function in the “lme4” package (Bates et al., 2015), and we rescaled and centered all continuous covariates at zero to achieve model convergence (Bolker et al., 2009). For LMMs, we used the “lme” function in the “nlme” package (Pinheiro et al., 2013).

## 3. RESULTS

### 3.1 Determining T_LETHAL_ via heat shock

Most embryos died at 45 or 46 °C (Fig. 2). There was no statistically clear effect of oviposition date (−0.006 ± 0.02 SE; t_1,69_ = −0.3; p=0.78) or initial egg mass (0.00004 ± 0.0003 SE; t_1,69_ = 0.01; p=0.99) on T_LETHAL_. Thus, we consider the intercept of the linear model, 45.3 °C (45.15 - 45.45; 95% CI), to be the mean value of T_LETHAL_.

**Figure 2.**
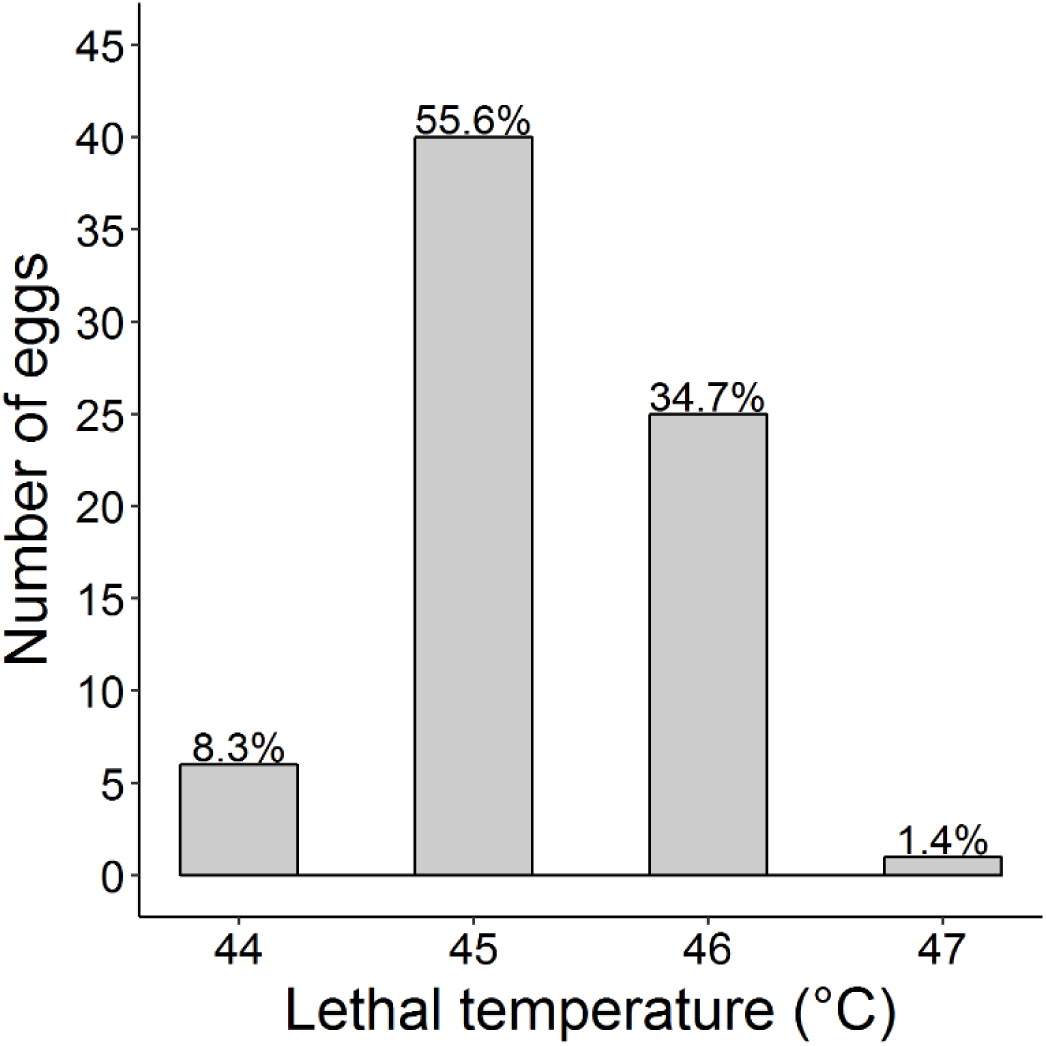
Histogram of T _LETHAL_ for*A. Sagrei* eggs exposed to 1-hour heat shocks. Values above each bar are the percentage of total eggs (n=72).

### 3.2 Thermal sensitivity of embryo physiology

Although the relationship between embryo heart rate and temperature was curvilinear (Table S1; Fig. 3), the linear component (3.40 ± 1.30 SE; df=97, t=2.62; p=0.01) of the regression was greater than the quadratic term (0.071 ± 0.019 SE; df=97, t=3.71; p=0.0003). The temperature at which heart rate was no longer detectable (i.e. T_LETHAL_) was 46.15 °C (45.87 - 46.44; 95% CI). This estimate of T_LETHAL_ differs from that of the heat shock experiment (i.e. confidence intervals do not overlap).

**Figure 3.**
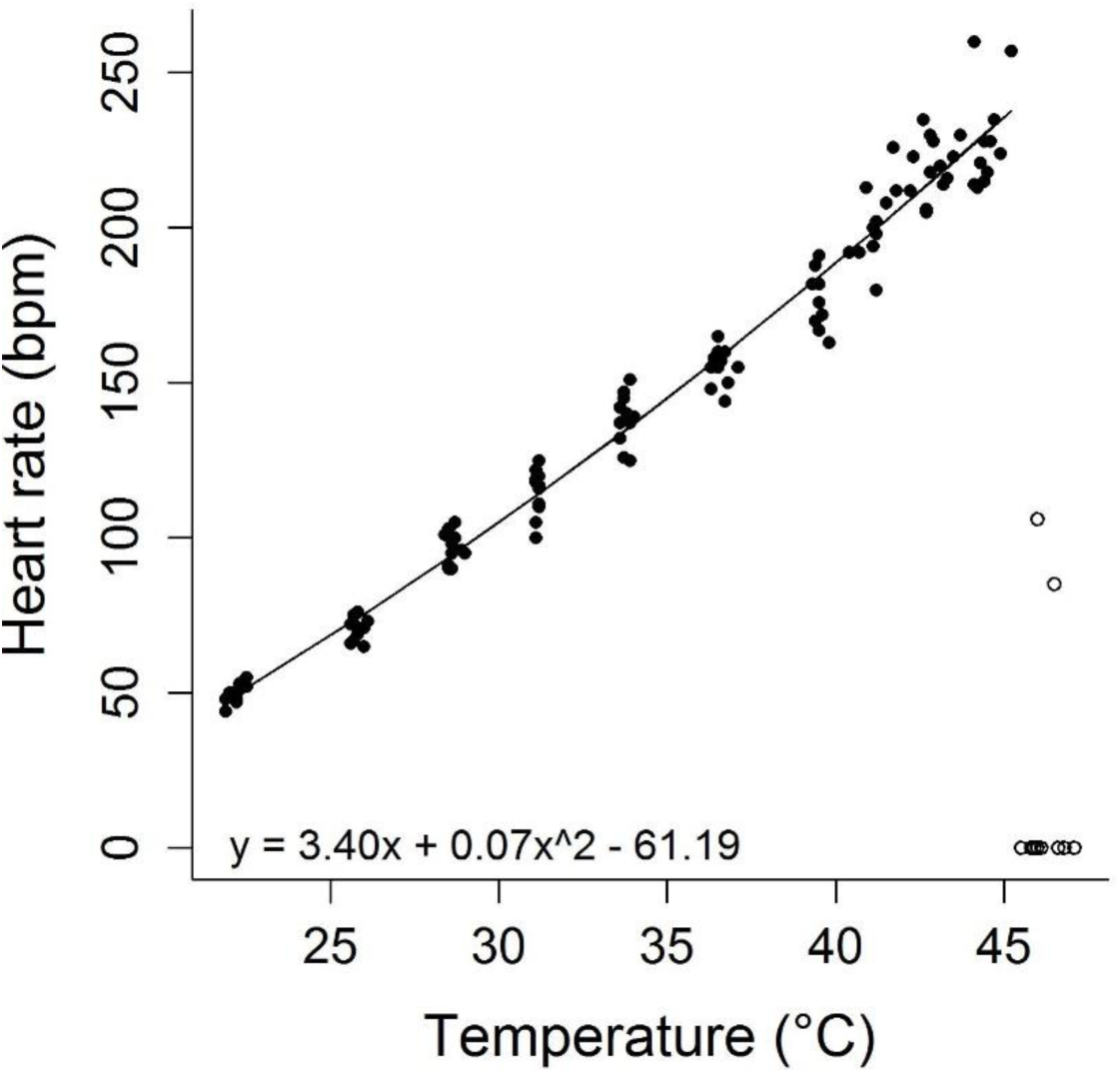
Heart rates of *A. sagrei* embryos across temperature. Closed and open circles show data that are include dor excluded from the regression, respectively.The equation for the regression is given in the panel.

VO_2_ of embryos was better explained by a second-degree polynomial than an asymptotic model (Table S1): VO_2_ increased steadily up to 40 °C, remained relatively constant from 39 to 42 °C and then declined (Fig. 4a). The linear component of VO_2_ by temperature was 3.82 (± 0.51 SE; df=37, t=7.43; p<0.0001) and the quadratic term was −0.046 (± 0.0078 SE; df=37, t=3.71; p<0.0001). RQ was stable from 22 to 39 °C, but steadily increased as embryos approached the lethal temperature (Fig. 4b). The rate of change in heart rate and VO_2_ for a 10 °C increase in temperature (i.e. Q_10_) was estimated using the equations in Fig 3 and Fig 4a, respectively. Q_10_s of heart rate and VO_2_ were similar at lower temperatures but diverged as embryos approached the lethal temperature (Fig. 5).

**Figure 4.**
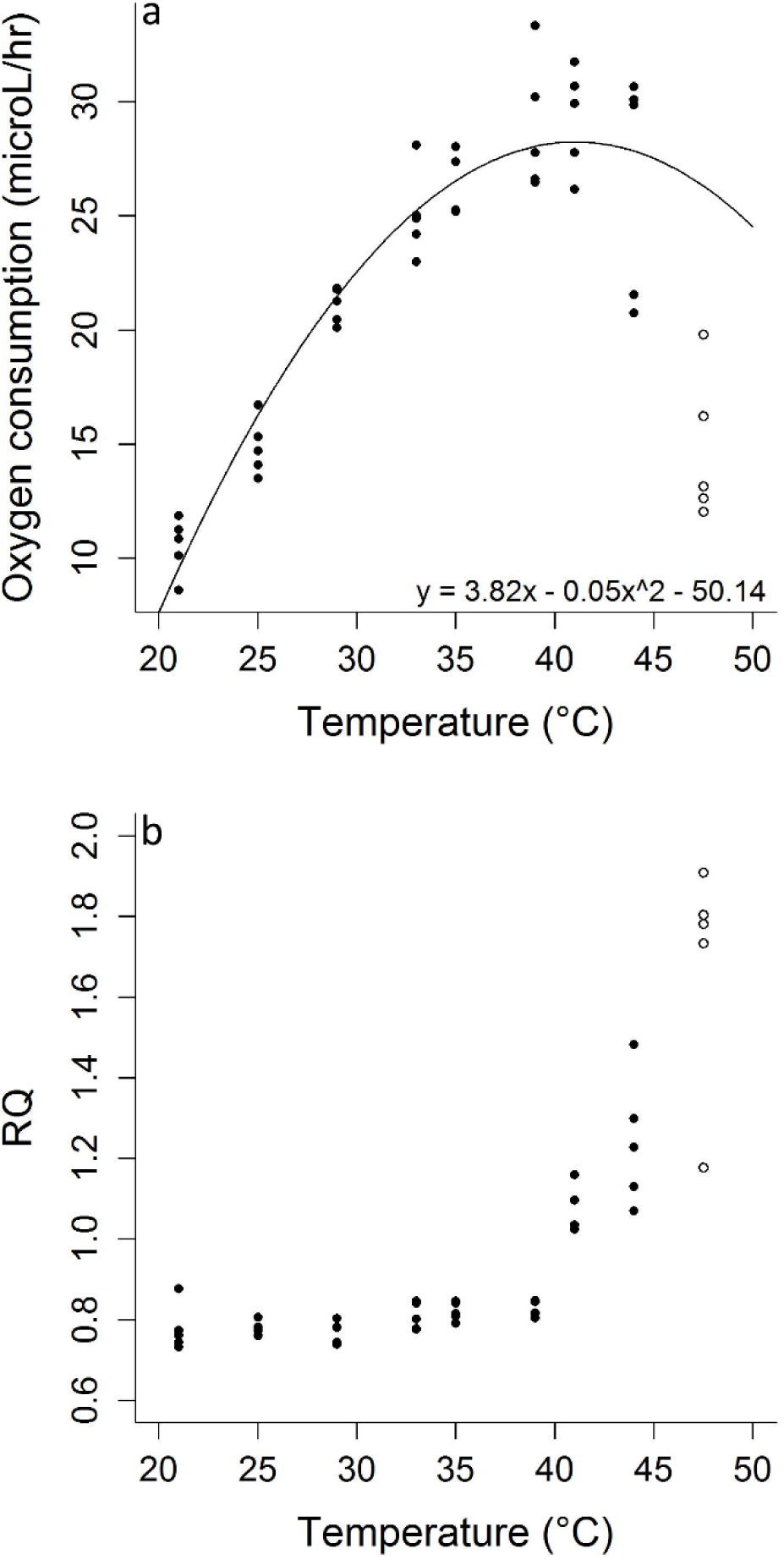
Oxygen consumption (a) and respiratory quotient (b)of *A. sagrei* embryos across temperature. Closed and open circles show raw data for eggs that did and did not survive measurements, respectively. Only surviving eggs were used to analyze oxygen consumption.The solid line in panel (a)shows the fit of a second-degree polynomial model.

**Figure 5.**
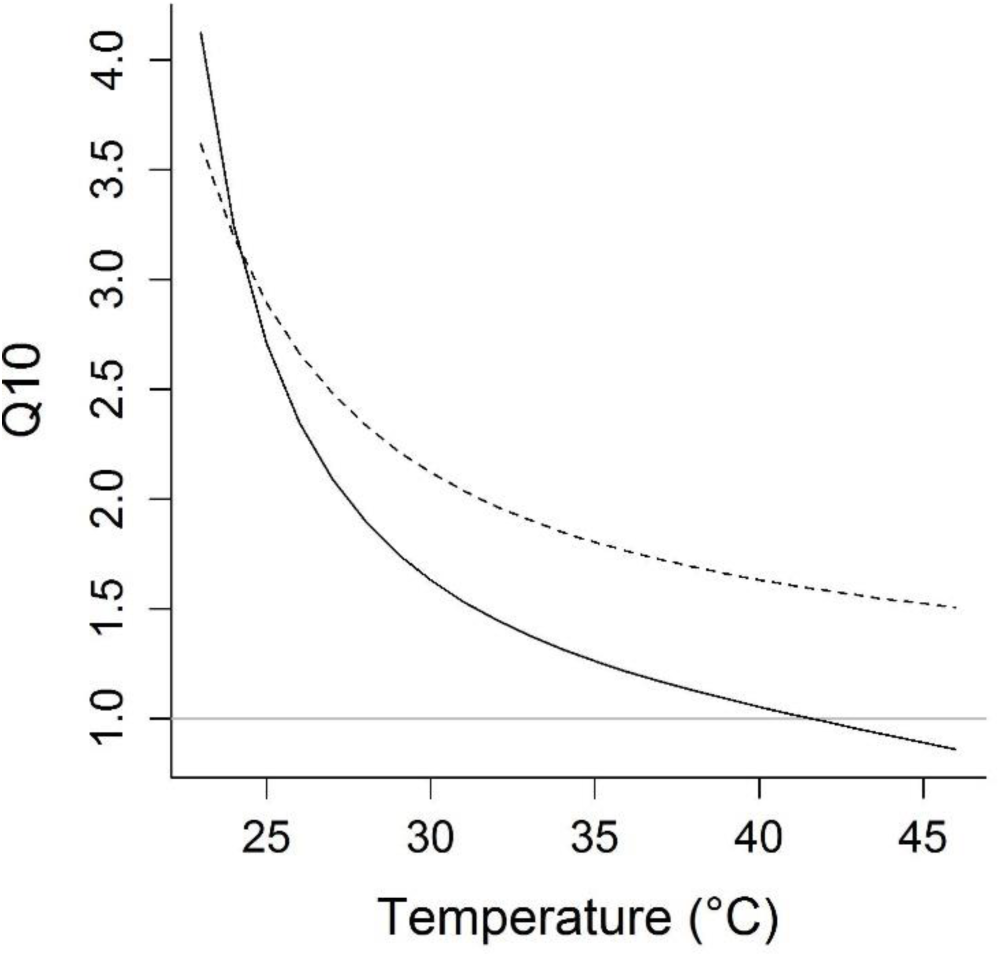
Temperature coefficient (i.e. Q_10_) of heart rate (dashed line) and oxygen consumption (solid line) across temperatures. A horizontal gray line shows the point at which reactions are insensitive to temperature (i.e. Q_10_= 1). Q_10_ values were calculated from estimates from the equations in Figures 3 and 4a. The Q_10_ shown at each temperature was calculated using that temperature and 1°C lower.

### 3.3 Repeated exposures to sublethal temperatures

Eggs exposed to four thermal spikes were 11.33 (± 2.96 SE) times as likely to die than controls (χ^21^=8.15; p=0.004). Raw mean survival was 97% and 75% for the control and experimental groups, respectively. This reduction in survival was largely influenced by embryo age at time of treatment (Table 1): for the experimental group, embryos were 1.42 (± 1.14 SE) times as likely to die with each 1 day increase in age (Fig. 6a). Thermal spikes reduced developmental rate by 0.025 (± 0.004 SE) stages per day (Table 1; Figure 6b). Given a mean rate of 0.504 stages per day for the control group, this equates to a 4.96% reduction.

**Table 1.** Results for final models testing the effects of treatment (0 or 4 thermal spikes at 43 °C peak temperature), embryo age at time of treatment and interactions and covariates on *A. sagrei* embryo and hatchling phenotypes and survival. Interactions with no statistics provided were omitted from the final model due to lack of statistical significance. See supplementary material for sample size, raw mean, and standard deviation of all response variables. For egg survival and developmental rates, initial mass was egg mass at oviposition. For hatchling morphology, growth, and survival, initial mass was hatchling mass at time of hatching. Results for egg survival are only for the experimental group (see statistical methods). Treatment estimates are the experimental group minus the control.

| Response variable | Initial mass |  | Age |  | Treatment |  |
| --- | --- | --- | --- | --- | --- | --- |
|  | Estimate (SE) |  | Estimate (SE) |  | Estimate (SE) |  |
| Egg survival** | 5.87 (29.44) | $\chi^2_1=0.04$; $p=0.84$ | -0.35 (0.13) | **$\chi^2_1=10.97$; $p=0.001$** | - | - |
| Developmental rate (stages/day) | -0.051 (0.14) | $F_{1,34}=0.13$; $p=0.72$ | 0.0001 (0.0005) | $F_{1,34}=0.05$; $p=0.82$ | -0.025 (0.0042) | **$F_{1,34}=35.05$; $p<0.0001$** |
| Hatchling initial SVL (mm) | 7.51 (5.00) | $F_{1,34}=2.3$; $p=0.14$ | 0.02 (0.02) | $F_{1,34}=1.2$; $p=0.29$ | -0.25 (0.18) | $F_{1,34}=2.0$; $p=0.16$ |
| Hatchling initial mass (g) | 0.36 (0.11) | **$F_{1,34}=11.0$; $p=0.002$** | 0.0006 (0.0004) | $F_{1,34}=2.5$; $p=0.13$ | -0.0080 (0.0032) | **$F_{1,34}=6.2$; $p=0.02$** |
| Hatchling initial tail length (mm) | 22.60 (12.00) | $F_{1,34}=3.6$; $p=0.07$ | 0.044 (0.039) | $F_{1,34}=1.3$; $p=0.27$ | -0.93 (0.34) | **$F_{1,34}=7.5$; $p=0.01$** |
| Hatchling survival | 0.44 (0.34) | $\chi^2_1=1.8$; $p=0.18$ | -0.46 (0.34) | $\chi^2_1=2.0$; $p=0.16$ | 1.31 (0.68) | **$\chi^2_1=4.0$; $p=0.045$** |
| Hatchling growth rate (g/day) | -0.024 (0.027) | $F_{1,20}=0.78$; $p=0.37$ | -0.00 (0.00) | $F_{1,20}=0.001$; $p=0.99$ | -0.0015 (0.00075) | $F_{1,20}=4.2$; $p=0.05$ |

**Figure 6.**
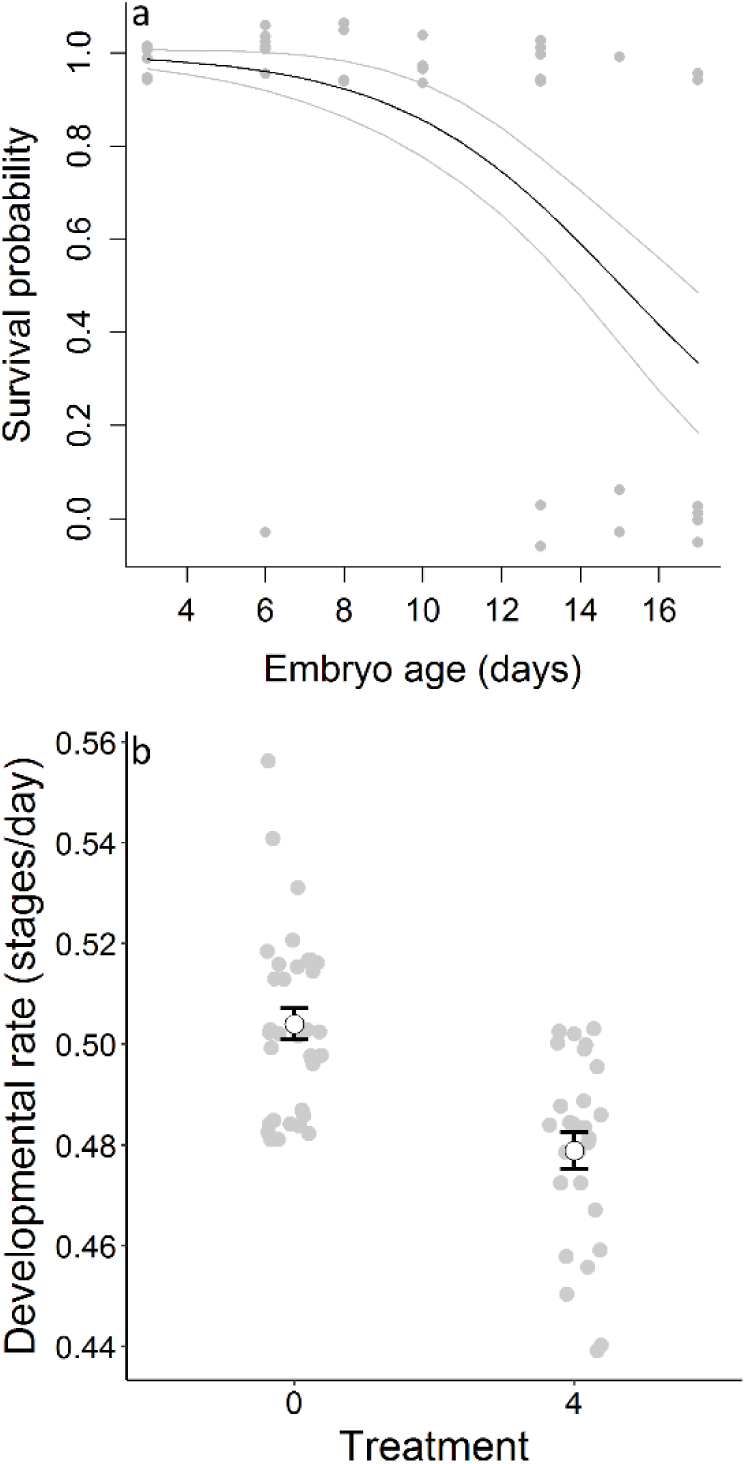
Effects of four exposures of thermal spikes on *A. sagrei* embryos. Panel a shows how survival probability of embryos in the experimental treatment declines with age. The solid line shows a regression of the raw data and gray circles are the raw data, jittered around 0 and 1 to avoid over-plotting. Panel b shows the effects of each treatment on developmental rate. Open circles are the raw mean, bars show standard error, gray circles show raw data.

We observed a statistically clear effect of the day by treatment interaction for embryo heart rate (F_2,123_ = 11.73; p<0.0001): heart rates of embryos exposed to four thermal spikes were 10.3 bpm (± 2.0 SE; p<0.0001) and 9.0 bpm (± 1.9 SE; p<0.0001) lower than controls on one and two days after experiencing the thermal spike, respectively (Fig. 7). These equate to 11.0 and 9.4 % reductions in heart rate, respectively.

**Figure 7.**
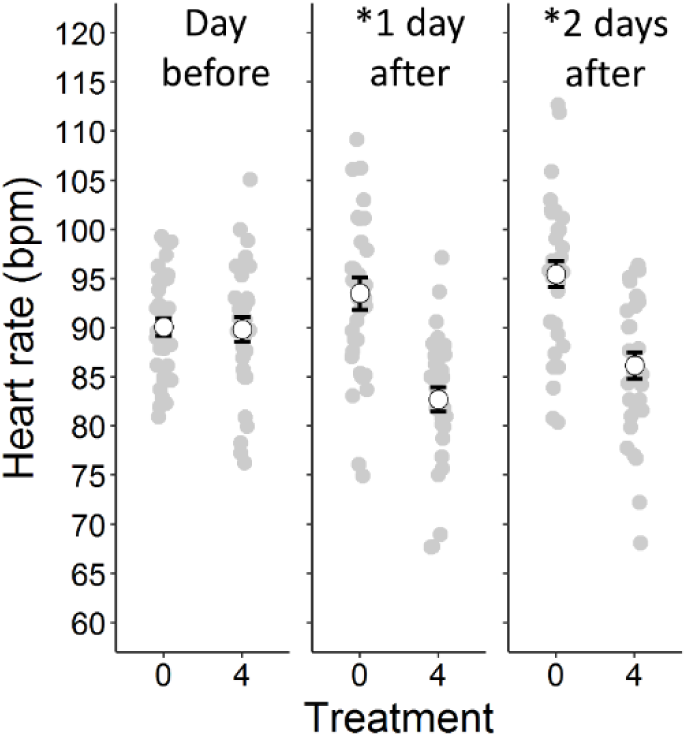
Heart rates of *A. sagrei* embryos exposed to 0 or 4 thermal spikes. Heart rates were measured at 28 °C on the day before exposure and on one and two days after exposure. Open circles show the raw means for each group, bars show standard error, and gray circles show the raw data. Asterisks signify a statistically significant difference in heart rate between groups.

Experimental hatchlings were shorter in SVL than those from the control, but this effect was not statistically clear (Table 1); however, experimental hatchlings were 0.01 (± 0.003 SE) g less massive than controls (Fig. 8a) and their tails were 0.93 mm (± 0.34 SE) shorter than controls (Fig. 8b; Table 1). This equates to a 5.9% reduction in body mass and a 3.1% reduction in tail length. Hatchlings exposed to thermal spikes were 3.72 (± 1.97 SE) times as likely to die as those from the control treatment (Fig. 8c; Table 1), and their growth rates were 0.0015 (± 0.00075 SE) g per day lower than controls (Fig. 8d). This equates to a 44% reduction in daily growth rate; this effect, however, was only marginally statistically clear (Table 1). See Supporting tables S2, S3, S4, and S5 for raw means, sample sizes, and standard deviations of all variables.

**Figure 8.**
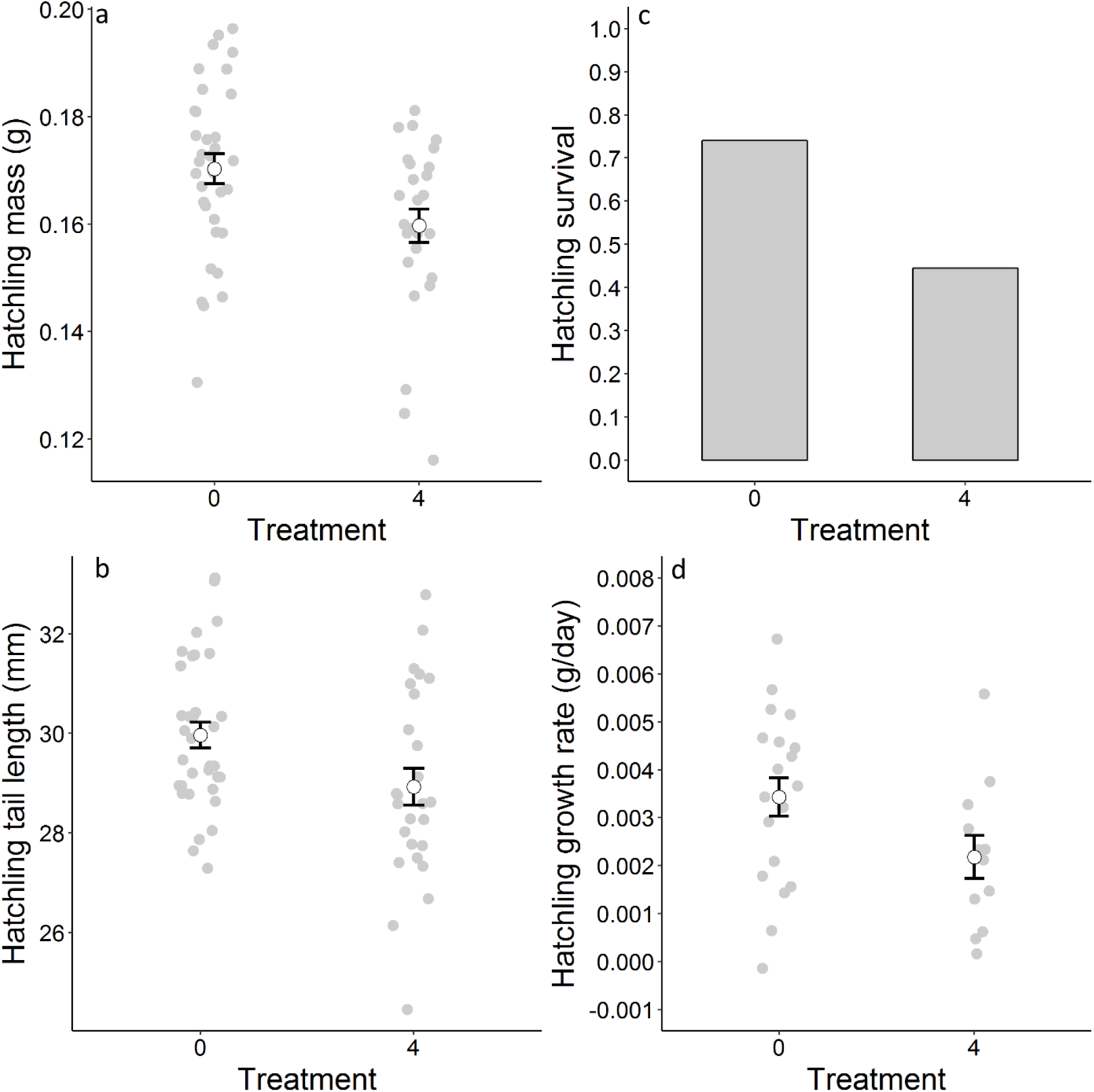
Effects of four exposures to extreme thermal fluctuations (43°C peak temperature) during development on hatchling mass (a) and tail length (b) at time of hatchling and hatchling survival (c) and growth rate (d) over a two-month period in the laboratory. In panels a, b, and d, open circles show the raw mean, bars show the standard error, and gray circles show the raw data.

## 4. DISCUSSION

Most studies of reptile embryo thermal tolerance utilize chronic, constant temperature treatments, and we know comparatively little about the effects of acute thermal stress on development. We subjected eggs of the brown anole lizard to brief exposures of high temperature to determine the T_LETHAL_ of embryos, quantify the thermal sensitivity of embryo physiology, and assess the cumulative effects of sub-lethal but stressful temperatures. Most embryos died at 45 or 46 °C, which is like that reported for other species (see below). Heart rate and VO_2_ increased across temperatures; however, as temperatures approached T_LETHAL_ heart rate and CO_2_ production increased while VO_2_ did not. Exposure to extreme fluctuations in nest temperature depressed developmental rates and embryo heart rates and resulted in hatchlings with smaller body size, reduced growth rates, and lower probability of survival. Thus, even brief exposure to extreme temperatures can have important effects on embryo development.

### 4.1 Determining T_LETHAL_ via heat shock

For reptile embryos, extreme (i.e. stressful) temperatures are generally considered to be those outside the range of constant temperatures that result in high hatching success (i.e. the optimal temperature range “OTR”; Andrews and Schwarzkopf 2012). The OTR has not been defined for *A. sagrei*, but it is probably between 20 and 33 °C. Eggs incubated from 26 to 30 °C have high hatching success (∼93%; Warner et al., 2011); however, Sanger et al., (2018) found that survival was high (90%) at 33 °C but low (39%) at 36 °C. In the latter study, eggs were dissected at day 12 and hatching success, *per se*, was not quantified. Given that thermal tolerance declines through development (see Fig. 6a), assessing egg survival via dissection, rather than hatching success, may overestimate the OTR. Other data (Pruett and Warner, unpublished) indicate that hatching success begins declining at 29°C and reaches zero at 35 °C. Thus, *A. sagrei* embryos can survive acute exposure to temperatures as much as 12 °C higher than the upper limit of the OTR. Thus, inferences about the thermal tolerance of species can be influenced by the use of chronic vs acute treatments. For example, embryos of the eastern fence lizard (*Sceloporus undulatus*) have adapted to pervasive (i.e. chronic) nest temperatures across a broad geographic range (Oufiero and Angilletta 2006; Du et al., 2010) such that northern populations develop more quickly than southern populations when incubated at a common temperature. However, embryo tolerance of acute thermal stress does not differ among populations (Angilletta et al., 2013). This is probably because even the most northerly populations experience nest temperatures above T_LETHAL_, potentially maximizing T_LETHAL_ across the range (Angilletta et al., 2013).

Although there are established protocols for quantifying the critical thermal maximum of adults, no such common protocols exist for estimating T_LETHAL_ of reptile embryos (Angilletta et al., 2013). We recommend that researchers consider the natural history and developmental ecology of their study species to determine the best method (e.g. chronic vs acute temperatures for deep vs shallow nesting species, respectively). For shallow-nesting species, like anoles, heat shock experiments may be appropriate; however, we did find some discrepancies among our estimates of T_LETHAL_. Our heart rate study, which utilized a thermal ramp, gave an estimate of T_LETHAL_ that was nearly 1 °C higher than the heat shock experiment (i.e. 46.2 vs 45.3 °C). Furthermore, a recent study that subjected eggs to thermal fluctuations of increasingly high temperature estimated T_LETHAL_ to be ∼ 44.5 °C for *A. sagrei* embryos (Hall and Warner, In press). Though each of these estimates were produced using ecologically meaningful treatments (i.e. acute exposure), the differences in T_LETHAL_ indicate that methods may influence estimates of thermal tolerance. This must be considered when designing experiments and making comparisons across the literature. One caveat is that we used different populations, which could account for variation in thermal sensitivity due to local adaptation (Du et al., 2010).

In reptiles (including birds), lethal temperature positively correlates with the optimal incubation temperature, which correlates with mean nest temperatures (Nechaeva 2011; Goa et al., 2014). This should generate a positive relationship between pervasive nest temperatures and thermal tolerance (Ma et al., 2018). Because *A. sagrei* nests can reach extremely warm temperatures (> 43 °C) during the hottest hours of the day (Sanger et al., 2018), we expect the upper thermal limit of *A. sagrei* embryos to be high compared to species that develop in cooler nests (e.g. *A. cristatellus*; Tiatragul et al., 2019; Hall and Warner, in review). However, few studies have determined the upper thermal limit of reptile embryos using acute exposures, which prevents broad comparisons among species. The few existing studies have found estimates of T_LETHAL_ similar to what we observed: embryos of the eastern fence lizard (*S. undulatus*), the plateau fence lizard (S. *tristichus*), the Chinese softshell turtle (*Pelodiscus sinensis*), and the Chinese glass lizard (*Takydromus septentrionalis*) die at 46, 45, 47, and 41 °C, respectively (Angilletta et al., 2013; Goa et al., 2014; Smith et al., 2015). More research is needed to understand how T_LETHAL_ varies across phylogeny and relates to ecological factors (e.g. optimal incubation temperature; mean nest temperatures). Given that many reptile species are threatened by climate change in part because mean temperatures and thermal variation of nests are increasing (Telemeco et al., 2009; Telemeco et al., 2017), a broadscale analysis of thermal tolerance of reptile embryos using acute exposures is warranted.

### 4.2 Thermal sensitivity of embryo physiology

The effect of temperature on embryo heart rate has been studied extensively in birds; however, much less attention has been given to non-avian reptiles (Nechaeva, 2011). Our results are consistent with existing studies. For example, Q_10_ of heart rate across nest temperatures was 2.18 for *A. sagrei* (Q_10_ of 2 to 3 for other reptiles; Du et al., 2011; Nechaeva, 2011). Q_10_ typically declines as temperature increases (Du et al., 2011) and we found this was true even up to the point of death (Fig. 5); however, at no point did heart rate become insensitive to temperature (i.e Q_10_ = 1). Q_10_ for VO_2_ and heart rate were similar at lower temperatures (23 to 30 °C); however, as temperatures approached T_LETHAL_, VO_2_ became less responsive to temperature compared with heart rate (Fig. 5).

To our knowledge, no other study has measured the thermal sensitivity of reptile embryo heart rate and VO_2_ across the full range of nest temperatures, including those near T_LETHAL_. At high temperatures, VO_2_ should plateau (i.e. maximum oxygen capacity, see Gangloff and Telemeco 2018), causing heart rate to decline or plateau due to low oxygen supply to cardiac muscle (Crossley and Altimiras 2005). Thus, we expected a similar relationship between heart rate and VO_2_ as temperatures approached T_LETHAL_. Rather, when approaching T_LETHAL_, heart rate continued rising. Moreover, RQ values increased near T_LETHAL_, indicating the use of anaerobic respiration since lizard embryos derive energy from lipids and protein (RQs between 0.7 and 0.9, Thompson et al., 2001). These data implicate a mismatch between oxygen demand and supply as a cause for death at high temperatures (oxygen and capacity limited thermal tolerance, Pörtner et al., 2017). Indeed, other studies find that hypoxic and hyperoxic incubation conditions decrease or increase, respectively, the thermal tolerance of reptile embryos (e.g. Smith et al., 2015; Liang et al., 2015); however, these have used chronic oxygen and/or temperature conditions. Thus, they demonstrate the positive correlation between oxygen supply and the critical thermal maximum but fail to unearth mechanisms that link the two. Our data indicate that death results from insufficient ventilation (i.e. diffusion of oxygen into the egg) rather than circulation (i.e. cardiac function) at high temperatures.

Contrary to our results, Angilletta et al. (2013) found that heart rates stabilized prior to death, which is the expected relationship for physiological performance curves (Angilletta 2006). The difference between their results and ours could be due to inter-specific variation in the thermal sensitivity of embryos. Indeed, reptile embryo heart rates vary widely according to phylogeny (Du et al., 2011). Moreover, due to ecological factors (e.g. shallow vs deep nests) embryo physiology may have adapted to respond to extreme temperatures in species-specific ways (Ma et al., 2018). Alternatively, we may not have measured heart rates across temperature at a fine enough scale to detect this stabilization phase. For example, heart rate substantially declined just prior to death for two individuals (open circles in Fig. 3), but we did not detect this for the other eggs. Regardless, our data indicate that the thermal sensitivity of heart rate and VO_2_ diverge at near-lethal temperatures, implicating a mismatch between oxygen supply and demand as a cause of death.

### 4.3 Repeated exposure to sublethal temperatures

In a separate study (Hall and Warner, Accepted), we subjected eggs to one or two thermal spikes with a peak of 43 °C. That study was conducted simultaneously with this one and utilized the same breeding colony, incubators, hatchling housing conditions, and incubation treatments. Therefore, results from these studies are comparable. For each response variable that was influenced by thermal spikes (egg survival, hatchling body size, hatchling growth and survival), the effect of 4 spikes was greater than that of 1 or 2. Thus, the negative effects of thermal spikes observed in this study represent an accumulation of damage rather than immediate effects of high temperature. Younger embryos were more robust to the treatment (Fig. 6a), possibly due to the relatively lower oxygen demand of early- vs late-stage embryos (Thompson and Stewart 1997). However, death at high temperatures likely results from multiple factors at different levels of biological complexity (Gangloff and Telemeco 2018) and additional research is required to understand how these effects combine to set the thermal limits of complex life.

Exposure to thermal spikes had substantial effects on physiology since both developmental rate and heart rate were suppressed by the treatment. Reduced developmental rates may have resulted directly from the reduction in heart rate since the total number of heart beats determines the incubation period in reptiles (Du et al., 2009). However, thermal spikes reduced developmental rates by 5%, but the observed decrease in heart rate (10% reduction for 2 days) can only account for 0.7% of this reduction. To fully account for the difference, heart rates would need to be depressed for the remainder of development, but we do not know how long this heart rate depression lasts. Depressed developmental rates can also result from diapause at extreme temperatures (Du et al., 2009); however, this is not likely since heart rate and VO_2_ remain relatively high across temperatures. Alternatively, high temperatures can induce cellular damage and subsequent repair (Sanger et al., 2018), which may slow developmental rates. Because hatchlings from the experimental treatment were smaller in body size, they may have diverted energy away from somatic growth and toward repair and maintenance.

The ecological implications of these data are difficult to assess. Slower developmental rates equate to longer incubation periods; thus, eggs are potentially exposed to adverse conditions for a longer time (e.g. egg depredation, extreme temperatures; Doody and Paul 2013). However, the effect we observed equates to a 1 or 2 day increase in the incubation period, which may not be biologically important. Hatchling body size and growth rates were reduced by thermal spikes, which may be responsible for the decreased rate of hatchling survival we observed. Indeed, larger hatchling body size can enhance survival probability (Sinervo et al., 1992); however, other factors, like the timing of hatching, may be much more important (Pearson and Warner 2018). Our study design (i.e. housing hatchlings communally) was ecologically meaningful since intraspecific competition is an important determinant of survival and growth for anoles (Calsbeek and Cox 2010); however, it prevents us from precisely identifying the cause of the detrimental effects on hatchlings. Reduced survival and growth may have resulted from the treatment *per se* (i.e. physiological effects) or from a diminished ability for experimental hatchlings to compete with those from the control group (i.e. ecological effects). Such effects could also combine or interact. A future study that measures performance and houses lizards individually and communally could assess such hypotheses.

The mean incubation temperatures for the control and experimental groups were 28.7 and 29.0 °C, respectively. If eggs were incubated at these two constant temperatures or at uniform, repeated fluctuations around them (e.g. sine wave), we would observe virtually no difference among treatments (Warner et al., 2011; Pearson and Warner 2016). Yet, the inclusion of a few extreme fluctuations resulted in significant depression of embryo physiology, reduced egg and hatchling survival, and reduced hatchling body size and growth. These data indicate that studies that solely utilize chronic incubation conditions poorly predict the effects of natural developmental environments. This is particularly true when nest temperatures fluctuate widely. In natural *A. sagrei* nests, temperatures often reach or exceed 43 °C (Sanger et al., 2018). How do embryos survive such extreme temperatures in the field? This is an open question; however, the answer may lie in the interactions between a myriad of abiotic factors that are present in the wild (e.g. moisture, solar radiation, oxygen availability, substrate composition, shade cover). The effects of such factors and their interactions with nest temperature on development are less often assessed than temperature alone (Warner et al., 2018) but are warranted.

### 4.4 Conclusions

Given the projected increases in the mean and variance of global temperatures, more research should be dedicated to understanding the effects of brief exposure of embryos to thermal stress (Burggren 2018). Indeed, when such exposures induce mortality, they may have more influence on the evolution of thermally sensitive traits than mean temperatures (Buckley and Huey 2016). Our results indicate that brown anole embryos have an upper lethal temperature that is much higher than the upper limit of the OTR. Moreover, we show that at near lethal temperatures there is a mismatch between oxygen demand and supply, which may contribute to death. However, when embryos are repeatedly exposed to temperatures below T_LETHAL_, thermal damage can accumulate, resulting in death or long-term effects on physiology that result in reduced survival of hatchlings. Thus, our study highlights the roles of both immediate and cumulative effects of high temperatures on embryo development.

## ACKNOWLEDGEMENTS

We thank A. Dees, A. DeSana, J Miracle, C. Reali, M. Turner, and K. Wilson for help with animal care, A. Appel for logistical advice, and C. Guyer for helpful comments on an earlier draft. This work was supported by the National Science Foundation (grant number DEB-1354897) and approved by the Auburn University IACUC committee (protocols 2017-3027, 2018-3233). This is publication number 907 of the Auburn University Museum of Natural history.

## Supporting information

### S1. Thermal sensitivity of metabolic rate for *A. sagrei* embryos

We used a Qubit Q-box RP1LP respirometer (Qubit Biology Inc., Kingston, ON) and applied a dynamic injection analysis method (see Lighton 2008, pgs 29-31). Prior to measurements, the length and width of each egg were recorded and used to estimate egg volume. This volume was later subtracted from the total volume of the respirometry chamber (i.e. a 10 ml Luer-Lok syringe) in order to calculate respiration rates. Each egg was placed in a 10 ml syringe with Luer-Lok tip attached to a 3-way stopcock (Becton, Dickinson and Company, Franklin Lakes, NJ 07417). A 5 µl drop of tap water was placed inside the syringe via a micropipette (to prevent desiccation). The plunger was inserted so the volume of the chamber was approximately 6 ml. The stopcock was hooked into the respirometer, and the syringe was flushed with CO_2_-free room air for 2 minutes at a rate of 100 ml/min. Two previously drilled holes (between the 5- and 6-ml marks) allowed air to exit the syringe (see Figure 4.3 in Lighton 2008). Prior to entering the syringe, this air was drawn through several meters of coiled tubing that was inside an incubator set to the target temperature for measurements. This ensured that the air used to flush the syringe was at the target temperature. A dummy syringe with a thermocouple inside of it was used to verify that the air temperature used for flushing was at the target temperature. After flushing, the syringe plunger was moved down to the 4 ml mark and the stop cock was twisted, sealing the egg in the syringe in a 4 ml volume of air (minus the volume of the egg and the drop of water). The time was noted, and eggs were placed in a constant temperature incubator set to the target temperature (either 21, 25, 29, 33, 35, 39, 41, 44, or 47 °C; n=5 eggs per temperature) for 30 minutes. At the end of 30 minutes, the syringe was removed from the incubator, a needle was affixed to the stopcock and a 2ml sample of air was injected into an injection port in the Qubit. The sample bolus was injected into a stream of dry, CO_2_-free air flowing at 50 ml/min. The time of injection was noted so we could calculate the exact time (in seconds) that each egg was respiring. Control syringes (containing a drop of water but no egg) were treated identically to those described previously and were injected into the respirometer at the beginning and end of the experiment.

The respirometer was calibrated prior to measurements for both CO_2_ and O_2_. For CO_2_, the zero value was calibrated by running room air through soda lime (to remove CO_2_). For O_2_, the zero was calibrated using O_2_-free gas (nitrogen). For CO_2_, we used outside air (assuming 400 ppm) to calibrate the higher end. For O_2_, we used dry (moisture removed via drierite) room air to calibrate the higher end at 20.95%.

**Figure S1.**
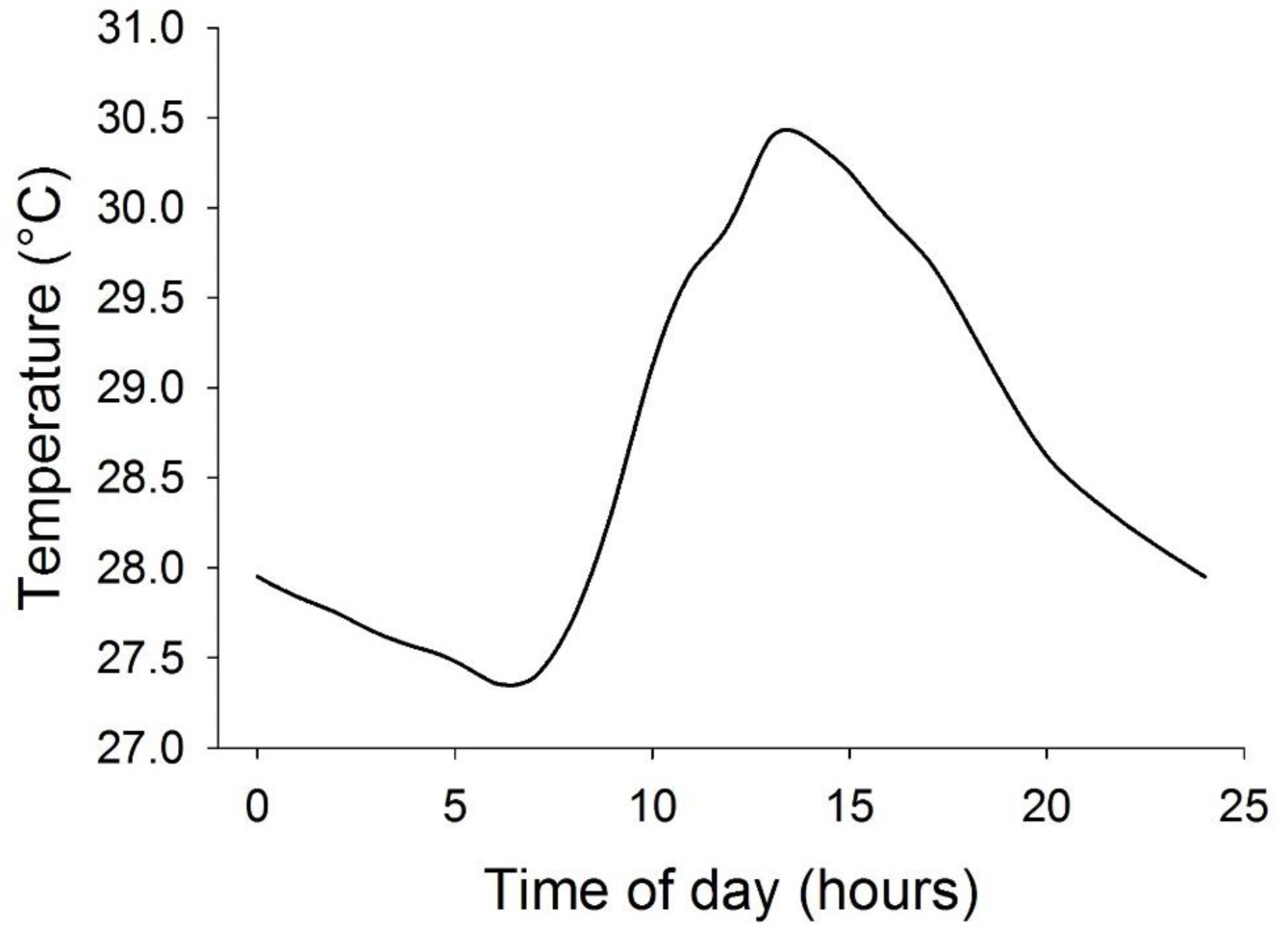
These temperatures are suitable for development of *A. sagrei* eggs (Tiatragul et al., 2017). Most eggs were incubated at these temperatures for the entirety of development. The only exception is that experimental eggs were removed from this fluctuation for 4 days to experience the thermal spikes. Control eggs were exposed to the fluctuation shown here during that time. Eggs used to determine the thermal sensitivity of heart rate and to measure rates of oxygen consumption were not incubated at these temperatures. They were incubated at a constant 28 °C until measurements of heart rate or metabolism were taken.

**Figure S2.**
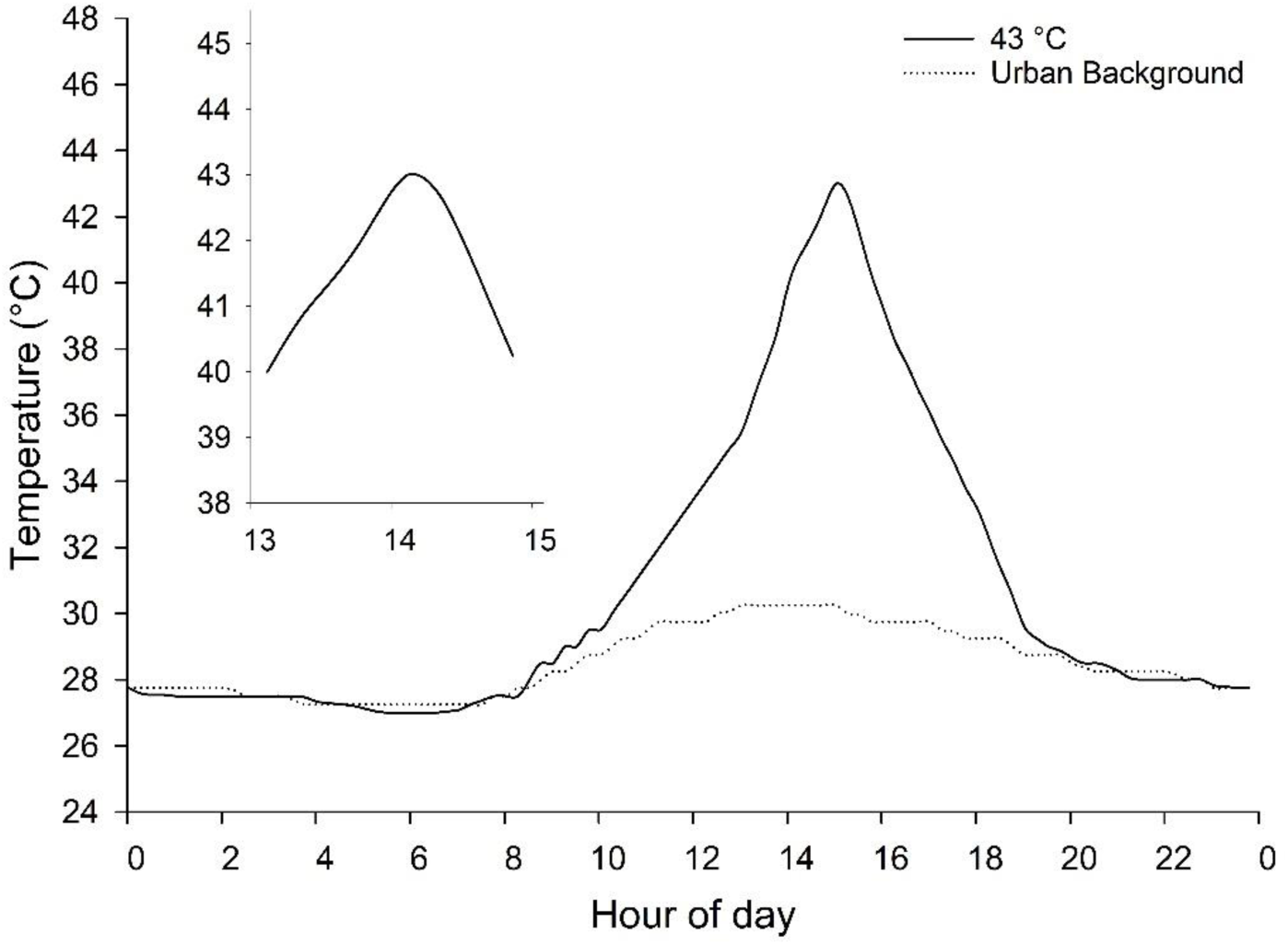
Temperature treatments used in our study. Fluctuations are based on field temperatures. The legend shows the peak temperature of the thermal spike. The inset figure in top left shows the peak at a finer scale. The urban background refers to the daily fluctuation used to incubate eggs before and after exposure to thermal spikes (i.e. Figure S1).

**Table S1.**
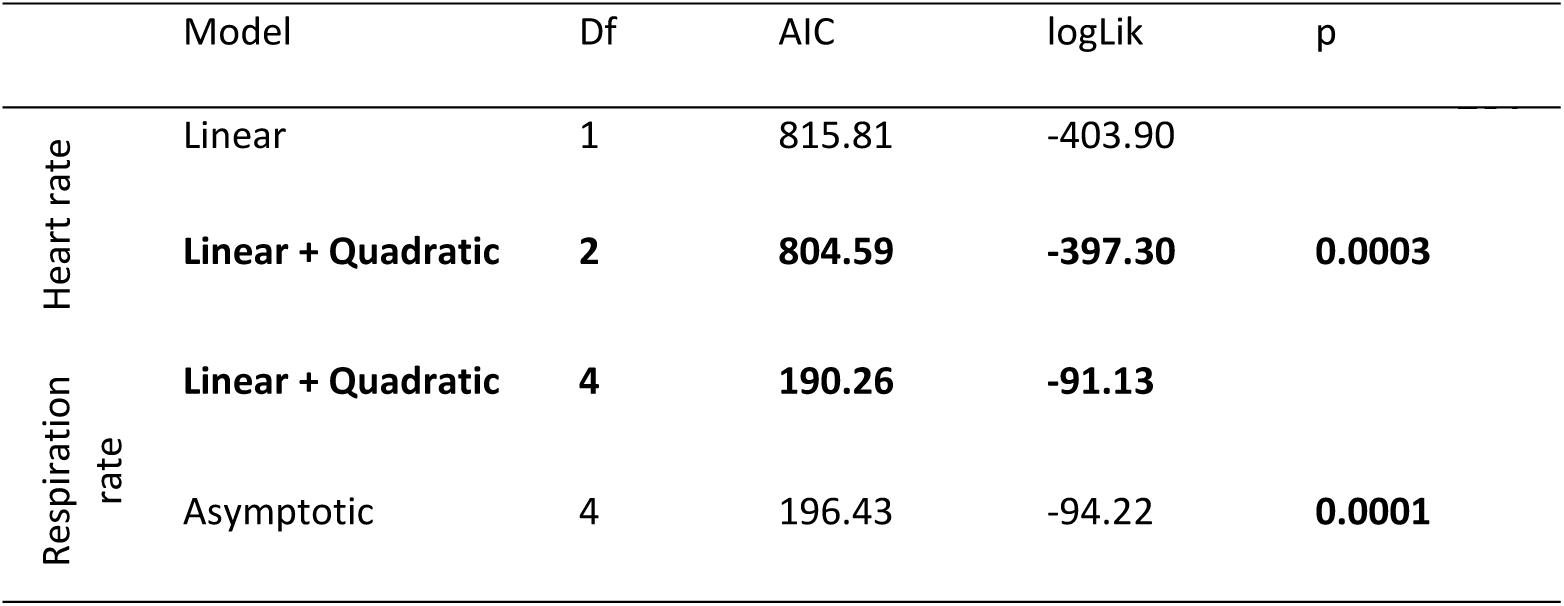
Comparison of linear and curvilinear relationship between *A. sagrei* embryo heart rate and temperature. Bold type denotes model chosen for analysis.

**Table S2.** Sample size (n), raw mean, and standard deviation (sd) for egg incubation period and hatchling phenotypes of *A. sagrei* exposed to one or four thermal spikes at 43 °C.

| Response variable | control |  |  | four spikes |  |  |
| --- | --- | --- | --- | --- | --- | --- |
|  | n | mean | sd | n | mean | sd |
| incubation period (days) | 34 | 29.79 | 1.04 | 27 | 31.37 | 1.24 |
| hatchling SVL (mm) | 34 | 18.04 | 0.73 | 27 | 17.73 | 0.6 |
| hatchling mass (g) | 34 | 0.1703 | 0.016 | 27 | 0.1597 | 0.0161 |
| hatchling tail length (mm) | 34 | 29.96 | 1.5 | 27 | 28.93 | 1.91 |
| hatchling growth rate (mg/day) | 20 | 3.43 | 1.78 | 12 | 2.18 | 1.55 |

**Table S3.** Sample sizes (n) and survival frequencies of eggs and hatchling lizards for *A. sagrei* exposed to zero (control) or four thermal spikes. Beneath each treatment (i.e. control, four spikes) the sample size for that treatment is given with the survival frequency for that sample in parenthesis.

| Egg survival |  |  | Hatchling survival |  |  |
| --- | --- | --- | --- | --- | --- |
| n | control | four spikes | n | control | four spikes |
| 71 | 35 (0.97) | 36 (0.75) | 53 | 26 (0.77) | 27 (0.44) |

**Table S4.**
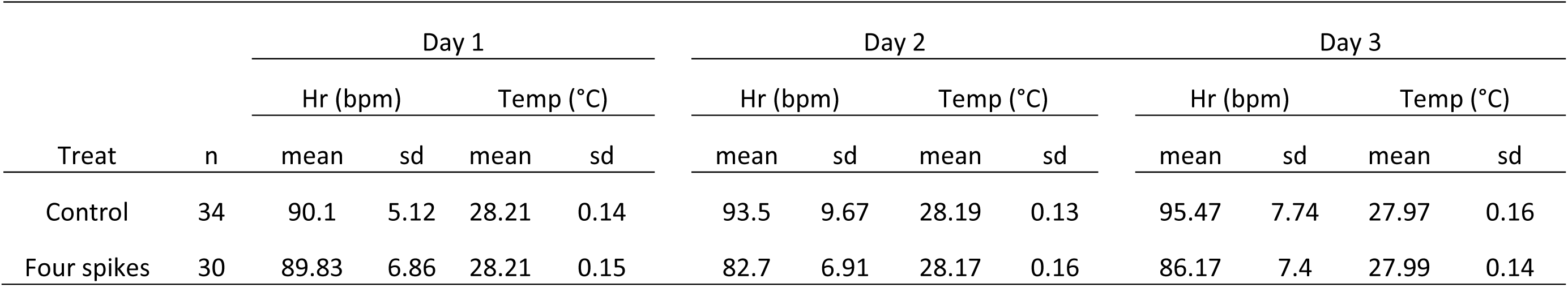
Mean and standard deviation (sd) for heart rates (Hr) and temperatures (Temp) at which heart rates were measured. Day 1 was the day before treatment (thermal spike), day 2 was the day after treatment, and day 3 was two days after treatment. Sample sizes (n) are listed for each treatment group. This was a repeated measures design and the same individuals were measured across all three days.

**S5.** Mean and standard deviation (sd) for the thermal sensitivity of anole heart rates (Hr) across nest temperatures.

**S5.** Mean and standard deviation (sd) for the thermal sensitivity of anole heart rates (Hr) across nest 2 temperatures.
| n | Hr (bpm) |  | Temp (°C) |  |
| --- | --- | --- | --- | --- |
|  | mean | sd | mean | sd |
| 11 | 50.00 | 3.10 | 22.22 | 0.23 |
| 11 | 70.73 | 3.58 | 25.79 | 0.17 |
| 11 | 96.73 | 5.18 | 28.64 | 0.18 |
| 11 | 114.82 | 7.55 | 31.15 | 0.05 |
| 11 | 138.27 | 8.19 | 33.76 | 0.14 |
| 11 | 155.18 | 5.98 | 36.58 | 0.24 |
| 11 | 176.78 | 9.60 | 39.50 | 0.14 |

## References

Andrews, R.M. and Schwarzkopf, L., 2012. Thermal performance of squamate embryos with respect to climate, adult life history, and phylogeny. Biological Journal of the Linnean Society, 106(4), pp. 851–864. https://doi.org/10.1111/j.1095-8312.2012.01901.x

Angilletta Jr, M.J., 2006. Estimating and comparing thermal performance curves. Journal of Thermal Biology, 31(7), pp. 541–545. https://doi.org/10.1016/j.jtherbio.2006.06.002

Angilletta, M.J., Zelic, M.H., Adrian, G.J., Hurliman, A.M. and Smith, C.D., 2013. Heat tolerance during embryonic development has not diverged among populations of a widespread species (Sceloporus undulatus). Conservation physiology, 1(1), pp. 1–9. https://doi.org/10.1093/conphys/cot018

Bates, D., Mächler, M., Bolker, B. and Walker, S., 2014. Fitting linear mixed-effects models using lme 4.67 (1) 1–51.

Battles, A.C. and Kolbe, J.J., 2019. Miami heat: Urban heat islands influence the thermal suitability of habitats for ectotherms. Global change biology, 25(2), pp. 562–576. https://doi.org/10.1111/gcb.14509

Bolker, B.M., Brooks, M.E., Clark, C.J., Geange, S.W., Poulsen, J.R., Stevens, M.H.H. and White, J.S.S., 2009. Generalized linear mixed models: a practical guide for ecology and evolution. Trends in ecology & evolution, 24(3), pp. 127–135. https://doi.org/10.1016/j.tree.2008.10.008

Booth, D.T., 2018. Incubation temperature induced phenotypic plasticity in oviparous reptiles: Where to next?. Journal of Experimental Zoology Part A: Ecological and Integrative Physiology, 329(6-7), pp. 343–350. https://doi.org/10.1002/jez.2195

Bowden, R.M., Carter, A.W. and Paitz, R.T., 2014. Constancy in an inconstant world: moving beyond constant temperatures in the study of reptilian incubation. Integrative and Comparative Biology, 54(5), pp. 830–840. https://doi.org/10.1093/icb/icu016

Buckley, L.B. and Huey, R.B., 2016. How extreme temperatures impact organisms and the evolution of their thermal tolerance. Integrative and comparative biology, 56(1), pp. 98–109. https://doi.org/10.1093/icb/icw004

Burggren, W., 2018. Developmental phenotypic plasticity helps bridge stochastic weather events associated with climate change. Journal of Experimental Biology, 221(9), p. jeb161984. https://doi.org/10.1242/jeb.161984

Calsbeek, R. and Cox, R.M., 2010. Experimentally assessing the relative importance of predation and competition as agents of selection. Nature, 465, pp. 613–616. https://doi.org/10.1038/nature09020

Carlo, M.A., Riddell, E.A., Levy, O. and Sears, M.W., 2018. Recurrent sublethal warming reduces embryonic survival, inhibits juvenile growth, and alters species distribution projections under climate change. Ecology letters, 21(1), pp. 104–116. https://doi.org/10.1111/ele.12877

Chalcraft, D.R. and Andrews, R.M., 1999. Predation on lizard eggs by ants: species interactions in a variable physical environment. Oecologia, 119(2), pp. 285–292. https://doi.org/10.1007/s004420050788

Cordero, G.A., Telemeco, R.S. and Gangloff, E.J., 2018. Reptile embryos are not capable of behavioral thermoregulation in the egg. Evolution & development, 20(1), pp. 40–47. https://doi.org/10.1111/ede.12244

Crossley, D.A. and Altimiras, J., 2005. Cardiovascular development in embryos of the American alligator *Alligator mississippiensis*: effects of chronic and acute hypoxia. Journal of Experimental Biology, 208(1), pp. 31–39. https://doi.org/10.1242/jeb.01355

Doody, J.S. and Paull, P., 2013. Hitting the ground running: environmentally cued hatching in a lizard. Copeia, 2013(1), pp. 160–165. https://doi.org/10.1643/CE-12-111

Du, W.G., Radder, R.S., Sun, B. and Shine, R., 2009. Determinants of incubation period: do reptilian embryos hatch after a fixed total number of heart beats?. Journal of Experimental Biology, 212(9), pp. 1302–1306. https://doi.org/10.1242/jeb.027425

Du, W.G., Warner, D.A., Langkilde, T., Robbins, T. and Shine, R., 2010. The physiological basis of geographic variation in rates of embryonic development within a widespread lizard species. The American Naturalist, 176(4), pp. 522–528. https://doi.org/10.1086/656270

Du, W.G., Ye, H., Zhao, B., Pizzatto, L., Ji, X. and Shine, R., 2011. Patterns of interspecific variation in the heart rates of embryonic reptiles. PloS one, 6(12), p. e29027. https://doi.org/10.1371/journal.pone.0029027

Du, W.G. and Shine, R., 2015. The behavioural and physiological strategies of bird and reptile embryos in response to unpredictable variation in nest temperature. Biological Reviews, 90(1), pp. 19–30. https://doi.org/10.1111/brv.12089

Gao, J., Zhang, W., Dang, W., Mou, Y., Gao, Y., Sun, B.J. and Du, W.G., 2014. Heat shock protein expression enhances heat tolerance of reptile embryos. Proceedings of the Royal Society B: Biological Sciences, 281(1791), pp. 1–10. https://doi.org/10.1098/rspb.2014.1135

Gangloff, E.J. and Telemeco, R.S., 2018. High temperature, oxygen, and performance: Insights from reptiles and amphibians. Integrative and comparative biology, 58(1), pp. 9–24. https://doi.org/10.1093/icb/icy005

Hall, J.M., Buckelew, A., Lovern, M., Secor, S.M. and Warner, D.A., 2018. Seasonal shifts in reproduction depend on prey availability for an income breeder. Physiological and Biochemical Zoology, 91(6), pp. 1129–1147. https://doi.org/10.1086/700341

Hall, J.M. and Warner, D.A., 2018. Thermal spikes from the urban heat island increase mortality and alter physiology of lizard embryos. Journal of Experimental Biology, 221(14), pp. 1–11. https://doi.org/10.1242/jeb.181552

Hall, J., Mitchell, T. and Warner, D., 2019. The brown anole (*Anolis sagrei*) as a model for studying life-history adaptation to seasonality. Anolis Newsletter. http://hdl.handle.net/11200/49424

Hall, J.M. and Warner D.A. In review. Thermal tolerance in the urban heat island: Thermal sensitivity varies ontogenetically and differs between embryos of two sympatric ectotherms. Journal of Experimental Biology.

Hulbert, A.C., Mitchell, T.S., Hall, J.M., Guiffre, C.M., Douglas, D.C. and Warner, D.A., 2017. The effects of incubation temperature and experimental design on heart rates of lizard embryos. Journal of Experimental Zoology Part A: Ecological and Integrative Physiology, 327(7), pp. 466–476. https://doi.org/10.1002/jez.2135

Kaiser, A., Merckx, T. and Van Dyck, H., 2016. The Urban Heat Island and its spatial scale dependent impact on survival and development in butterflies of different thermal sensitivity. Ecology and Evolution, 6(12), pp. 4129–4140. https://doi.org/10.1002/ece3.2166

Liang, L., Sun, B.J., Ma, L. and Du, W.G., 2015. Oxygen-dependent heat tolerance and developmental plasticity in turtle embryos. Journal of Comparative Physiology B, 185(2), pp. 257–263. https://doi.org/10.1007/s00360-014-0874-4

Ma, L., Sun, B.J., Li, S.R., Hao, X., Bi, J.H. and Du, W.G., 2018. The vulnerability of developing embryos to simulated climate warming differs between sympatric desert lizards. Journal of Experimental Zoology Part A: Ecological and Integrative Physiology, 329(4-5), pp. 252–261. https://doi.org/10.1002/jez.2179

McDonald, R.I., Kareiva, P. and Forman, R.T., 2008. The implications of current and future urbanization for global protected areas and biodiversity conservation. Biological conservation, 141(6), pp. 1695–1703. https://doi.org/10.1016/j.biocon.2008.04.025

Nechaeva, M.V., 2011. Physiological responses to acute changes in temperature and oxygenation in bird and reptile embryos. Respiratory physiology & neurobiology, 178(1), pp. 108–117. https://doi.org/10.1016/j.resp.2011.04.003

Noble, D.W., Stenhouse, V. and Schwanz, L.E., 2018. Developmental temperatures and phenotypic plasticity in reptiles: A systematic review and meta-analysis. Biological Reviews, 93(1), pp. 72–97. https://doi.org/10.1111/brv.12333

Oufiero, C.E. and Angilletta Jr, M.J., 2006. Convergent evolution of embryonic growth and development in the eastern fence lizard (*Sceloporus undulatus*). Evolution, 60(5), pp. 1066–1075. https://doi.org/10.1111/j.0014-3820.2006.tb01183.x

Pearson, P.R. and Warner, D.A., 2016. Habitat-and season-specific temperatures affect phenotypic development of hatchling lizards. Biology letters, 12(10), pp. 1–5. https://doi.org/10.1098/rsbl.2016.0646

Pearson, P.R. and Warner, D.A., 2018. Early hatching enhances survival despite beneficial phenotypic effects of late-season developmental environments. Proceedings of the Royal Society B: Biological Sciences, 285(1874), pp. 1–9. https://doi.org/10.1098/rspb.2018.0256

Pinheiro, J., Bates, D., DebRoy, S. and Sarkar, D., 2013. the R Development Core Team (2013) nlme: linear and nonlinear mixed effects models. R package version, 3, pp. 1–97.

Pörtner, H.O., Bock, C. and Mark, F.C., 2017. Oxygen-and capacity-limited thermal tolerance: bridging ecology and physiology. Journal of Experimental Biology, 220(15), pp. 2685–2696. https://doi.org/10.1242/jeb.134585

R Core Team. 2018. R: A language and environment for statistical computing. R Foundation for Statistical Computing, Vienna, Austria. http://www.R-project.org/.

Refsnider, J.M., Clifton, I.T. and Vazquez, T.K., 2019. Developmental plasticity of thermal ecology traits in reptiles: Trends, potential benefits, and research needs. Journal of Thermal Biology, 84, pp. 74–82. https://doi.org/10.1016/j.jtherbio.2019.06.005

Sanger, T.J., Hime, P.M., Johnson, M.A., Diani, J. and Losos, J.B., 2008a. Laboratory protocols for husbandry and embryo collection of *Anolis* lizards. Herpetological Review, 39(1), pp. 58–63.

Sanger, T.J., Losos, J.B. and Gibson-Brown, J.J., 2008b. A developmental staging series for the lizard genus *Anolis*: a new system for the integration of evolution, development, and ecology. Journal of Morphology, 269(2), pp. 129–137. https://doi.org/10.1002/jmor.10563

Sanger, T.J., Kyrkos, J., Lachance, D.J., Czesny, B. and Stroud, J.T., 2018. The effects of thermal stress on the early development of the lizard *Anolis sagrei*. Journal of Experimental Zoology Part A: Ecological and Integrative Physiology, 329(4-5), pp. 244–251. https://doi.org/10.1002/jez.2185

Santidrián Tomillo, P., Genovart, M., Paladino, F.V., Spotila, J.R. and Oro, D., 2015. Climate change overruns resilience conferred by temperature-dependent sex determination in sea turtles and threatens their survival. Global change biology, 21(8), pp. 2980–2988. https://doi.org/10.1111/gcb.12918

Shine, R., Elphick, M.J. and Barrott, E.G., 2003. Sunny side up: lethally high, not low, nest temperatures may prevent oviparous reptiles from reproducing at high elevations. Biological Journal of the Linnean Society, 78(3), pp. 325–334. https://doi.org/10.1046/j.1095-8312.2003.00140.x

Shine, R., Langkilde, T., Wall, M. and Mason, R.T., 2005. The fitness correlates of scalation asymmetry in garter snakes *Thamnophis sirtalis parietalis*. Functional Ecology, 19(2), pp. 306–314. https://doi.org/10.1111/j.1365-2435.2005.00963.x

Sinervo, B., Zamudio, K., Doughty, P. and Huey, R.B., 1992. Allometric engineering: a causal analysis of natural selection on offspring size. Science, 258(5090), pp. 1927–1930. https://doi.org/10.1126/science.258.5090.1927

Sinervo, B., Mendez-De-La-Cruz, F., Miles, D.B., Heulin, B., Bastiaans, E., Villagrán-Santa Cruz, M., Lara-Resendiz, R., Martínez-Méndez, N., Calderón-Espinosa, M.L., Meza-Lázaro, R.N. and Gadsden, H., 2010. Erosion of lizard diversity by climate change and altered thermal niches. Science, 328(5980), pp. 894–899. https://doi.org/10.1126/science.1184695

Smith, C., Telemeco, R.S., Angilletta Jr, M.J. and VandenBrooks, J.M., 2015. Oxygen supply limits the heat tolerance of lizard embryos. Biology letters, 11(4), pp. 1–4. https://doi.org/10.1098/rsbl.2015.0113

Telemeco, R.S., Elphick, M.J. and Shine, R., 2009. Nesting lizards (*Bassiana duperreyi*) compensate partly, but not completely, for climate change. Ecology, 90(1), pp. 17–22. https://doi.org/10.1890/08-1452.1

Telemeco, R.S., Fletcher, B., Levy, O., Riley, A., Rodriguez-Sanchez, Y., Smith, C., Teague, C., Waters, A., Angilletta Jr, M.J. and Buckley, L.B., 2017. Lizards fail to plastically adjust nesting behavior or thermal tolerance as needed to buffer populations from climate warming. Global change biology, 23(3), pp. 1075–1084. https://doi.org/10.1111/gcb.13476

Tiatragul, S., Kurniawan, A., Kolbe, J.J. and Warner, D.A., 2017. Embryos of non-native anoles are robust to urban thermal environments. Journal of thermal biology, 65, pp. 119–124. https://doi.org/10.1016/j.jtherbio.2017.02.021

Tiatragul, S., Hall, J.M., Pavlik, N.G. and Warner, D.A., 2019. Lizard nest environments differ between suburban and forest habitats. Biological Journal of the Linnean Society, 126(3), pp. 392–403. https://doi.org/10.1093/biolinnean/bly204

Tiatragul, S., Hall J.M., Warner D.A. In review. Nestled in the city heat: nesting behavior facilitates embryo development in an urbanised environment. Journal of Urban Ecology

Thompson, M.B. and Stewart, J.R., 1997. Embryonic metabolism and growth in lizards of the genus *Eumeces*. Comparative Biochemistry and Physiology Part A: Physiology, 118(3), pp. 647–654. https://doi.org/10.1016/S0300-9629(97)00081-9

Thompson, M.B., Speake, B.K., Russell, K.J. and McCartney, R.J., 2001. Utilisation of lipids, protein, ions and energy during embryonic development of Australian oviparous skinks in the genus *Lampropholis*. Comparative Biochemistry and Physiology Part A: Molecular & Integrative Physiology, 129(2-3), pp.313–326. https://doi.org/10.1016/S1095-6433(00)00349-4

Warner, D.A., Moody, M.A., Telemeco, R.S. and Kolbe, J.J., 2011. Egg environments have large effects on embryonic development, but have minimal consequences for hatchling phenotypes in an invasive lizard. Biological Journal of the Linnean Society, 105(1), pp. 25–41. https://doi.org/10.1111/j.1095-8312.2011.01778.x

Warner, D.A. and Shine, R., 2011. Interactions among thermal parameters determine offspring sex under temperature-dependent sex determination. Proceedings of the Royal Society B: Biological Sciences, 278(1703), pp. 256–265. https://doi.org/10.1098/rspb.2010.1040

Warner, D.A., Du, W.G. and Georges, A., 2018. Introduction to the special issue-Developmental plasticity in reptiles: Physiological mechanisms and ecological consequences. Journal of experimental zoology. Part A, Ecological and integrative physiology, 329(4-5), pp.153–161. https://doi.org/10.1002/jez.2199

While, G.M., Noble, D.W., Uller, T., Warner, D.A., Riley, J.L., Du, W.G. and Schwanz, L.E., 2018. Patterns of developmental plasticity in response to incubation temperature in reptiles. Journal of Experimental Zoology Part A: Ecological and Integrative Physiology, 329(4-5), pp.162–176. https://doi.org/10.1002/jez.2181

